# Protein structure dynamic prediction: a Machine Learning/Molecular Dynamic approach to investigate the protein conformational sampling

**DOI:** 10.1101/2021.12.01.470731

**Authors:** Martina Audagnotto, Werngard Czechtizky, Leonardo De Maria, Helena Käck, Garegin Papoian, Lars Tornberg, Christian Tyrchan, Johan Ulander

## Abstract

Proteins exist in several different conformations. These structural changes are often associated with fluctuations at the residue level. Recent findings show that co-evolutionary analysis coupled with machine-learning techniques improves the precision by providing quantitative distance predictions between pairs of residues. The predicted statistical distance distribution from Multi Sequence Analysis (MSA) reveals the presence of different local maxima suggesting the flexibility of key residue pairs. Here we investigate the ability of the residue-residue distance prediction to provide insights into the protein conformational ensemble. We combine deep learning approaches with mechanistic modeling to a set of proteins that experimentally showed conformational changes. The predicted protein models were filtered based on energy scores, RMSD clustering, and the centroids selected as the lowest energy structure per cluster. The models were compared to the experimental-Molecular Dynamics (MD) relaxed structure by analyzing the backbone residue torsional distribution and the sidechain orientations. Our pipeline not only allows us to retrieve the global experimental folding but also the experimental structural dynamics. We show the potential correlation between the experimental structure dynamics and the predicted model ensemble demonstrating the susceptibility of the current state-of-the-art methods in protein folding and dynamics prediction and pointing out the areas of improvement.

## Introduction

Understanding the relationship between protein 3D structure and function is the key to unveil the protein biological mechanism and therefore the ability to modulate it. The Anfinsen’s thermodynamic hypothesis states that all the information driving the protein folding is encoded in the protein first sequence.^1, 2^ From a kinetic perspective, there is an almost linear correlation (R=0.75) between the natural logarithm of folding rate and the percentage of Relative Contact Order (%RCO)^3^ which shows the dependency of the kinetic folding from the overall protein topology. According to the Levinthal paradox^4^, despite the huge number of potential conformations, a protein can fold to its one conformation, namely native structure, following folding pathways characterized by folding intermediates^5, 6^. Studies of the polymer chain entropies revealed that low-energy intermediate conformational ensembles are less populated compared to high-energy ensembles indicating that the protein-folding energy landscapes are funnel-shaped^5, 7–10^. The funneled formulation offers insights into proteins conformational heterogeneity and protein chains entropy providing a microscopic framework for folding kinetics.

There are two main computational approaches used to predict 3D protein structures: (i) the template-based modeling (TBM) and (ii) the free-modeling approaches (FM). TBM is based on the observation that homologous proteins have similar structure so a known protein structure can be used as template for homologous protein^11^. This approach is limited by the heterogeneity of the experimentally available structures in the PDB which, up to date, count only 1200 different folds^12^. The FM approach, also known as *ab initio* prediction, is required for proteins that lack any statistically significant similar protein sequences with known structures. Among the plethora of possible FM approaches extensively used ones are the fragment assembly ^13–16^ or the first principle physics-based methods^17–19^. These methods are based on physics-based force fields to search for low energy states following the Anfinsen’s hypothesis. However, they suffer from the inaccuracy of the force fields potential in the description of the thermodynamic protein stability which lead to an inaccurate description of the atomic interaction and folds.

The application of deep learning methods to the protein structure prediction field is a game changer. In 2019, during the CASP13 competition, several groups (Rosetta, DeepMind and FEIG-R2 just to name a few) showed that by applying deep learning based methods it was possible to gain fold-level accuracy for proteins lacking homologues in the PDB. By combining deep learning approaches with inputs from coevolutionary coupling features derived from Multi Sequence Alignment (MSA), the level of accuracy (GDT-TS) reached was ≈ 10% higher (for the easy targets) and ≈ 20% higher (for the difficult targets) than in the previous version of CASP^20^. Indeed, these methods showed that evolutionary information captured in MSA of homologous sequences not only offers the template of known structure but also inter-residue correlations that can be decoded into contact maps. These contact maps are then translated into distograms (distance probability distribution plots) providing quantitative information that can be used as potential restraints to improve the accuracy of *de novo* structure prediction of static protein structure.

The impressive performance of AlphaFold2^21^ in CASP14 demonstrated that deep learning techniques have reached a point where the accuracy of the predicted model, in many cases, is comparable with that of the experimental structure. With this new generation of computational structure prediction tools we can focus on many other questions with the highest biological relevance such as (i) small-molecule docking to protein models, (ii) protein-protein interaction predictions and (iii) protein conformational ensemble prediction. The latter is the focus of the discussion reported in this article.

.Multiple conformations are often linked to different protein functions, e.g. active and inactive conformation in enzymes, and inward versus outward facing upon substrate transport binding in transporters. It has been long shown that in the PDB ^12^ one can find entries with similar or even identical sequences that have different structures. Since, deep learning methods can be trained over the entire PDB and it has been proven that MSA can encode information about functional conformations^22^, it is reasonable to hypothesize that the current prediction methods would be able to propose biological relevant conformation of a query protein. In CASP14, AlphaFold2 predicted the accurate structure of 75 targets within the submitted 5 models. Masrati et al.^23^ superimposed and visually inspected all the 5 models for each of the 75 predicted targets finding that 80% showed the same conformations while the remaining 20% have more than one single distinct one.

To test the ability of exploring the conformational space of a query protein we built up a pipeline with trRosetta^24^ as a core. By combining the current state-of-the-art ML technique for MSA (deepMSA) with deep residual-convolutional network transform-restrain Rosetta (trRosetta) and the AWSEM force field, we observed that not only the X-ray structures of the different protein states were predicted but also the similar intermediate states explored by MD simulation run on the experimental structures.

Currently, it is difficult to assess the ability of AlphaFold2 in predicting multiple biologically relevant conformations for several different proteins. Although the code is available, it requires expensive hardware and a storage data capability of five TB as reported in the alphafold github (https://github.com/deepmind/alphafold). A Jupyter Notebook version, namely ColabFold^25^, was recently developed. It combines AlphaFold2 with Google Colab^25^ providing a free and accessible on-line platform for protein folding that does not require any installation. We therefore decided to adopt the ColabFold pipeline to generate five models and to compare them with the predicted structural ensembles generated by our pipeline.

## Results

### Test case database

To test the ability of our pipeline to predict protein conformational ensembles, we investigated only X-ray structures with a maximum length of roughly 200 amino acids, a resolution equal or less than 2.40 Å and where more than one conformation was available in the PDB for the same sequence. Our analysis set is composed of a total of 5 structures: 4 proteins from the Conformational Diversity in the Native State database (CoDNas)^26^, 1 protein from the CASP14 list. These structures represent examples of conformational changes in proteins induced by small molecules, metal coordination ion or point mutation. Each of the test cases has the peculiarity of having multiple PDB structures for single protein sequences. For our analysis the pairs with maximum RMSD between conformers have been considered.. The PDB structures were preprocessed to remove small molecules and ions. The list of PDB structures is summarized in Table 1.

**Table 1.**
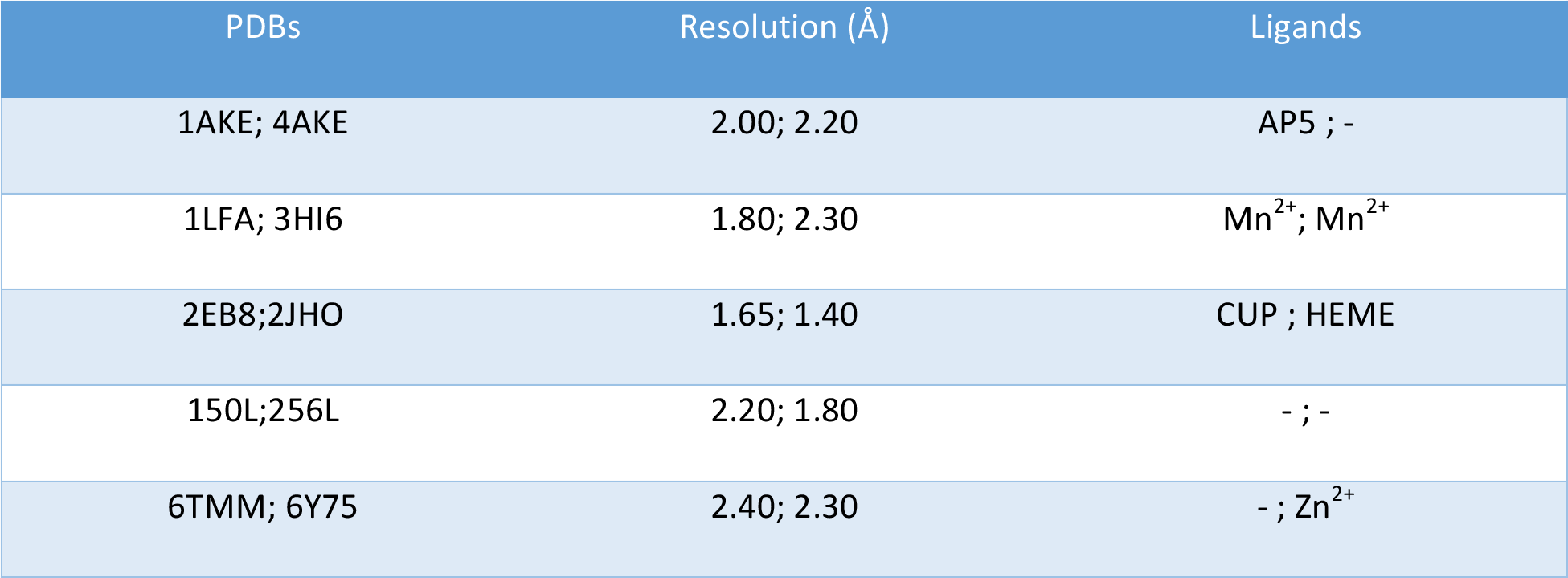
Test cases list.

| PDBs | Resolution (Å) | Ligands |
| --- | --- | --- |
| 1AKE; 4AKE | 2.00; 2.20 | AP5 ; - |
| 1LFA; 3HI6 | 1.80; 2.30 | Mn<sup>2+</sup>; Mn<sup>2+</sup> |
| 2EB8;2JHO | 1.65; 1.40 | CUP ; HEME |
| 150L;256L | 2.20; 1.80 | - ; - |
| 6TMM; 6Y75 | 2.40; 2.30 | - ; Zn<sup>2+</sup> |

### Adenylate kinase (adk)

Adenylate kinase isolated from Escherichia coli (AK_e_) is a small phosphotransferase enzyme in which the conversion between active (open) and inactive (closed) conformations is rate limiting for catalysis. AK_e_ enzyme consists of a CORE subdomain (from N79 to G214 residue), AMP- (from V121 to Q160 residue) and ATP- (from I26 to R78 residue) binding subdomains which undergo significant conformational changes upon substrate binding. This enzyme catalyzes a reversible phosphoryl transfer reaction with the primary purpose of maintaining the energy balance in cells, and is ubiquitously expressed in many different organisms^27–29^. From a functional point of view, the conformational shift causes the rearrangement of the backbone H-bonding patterns without unfolding the entire protein.

The structure of AK_e_ has been solved for substrate-free (open, PDB:1AKE)^28^ and inhibitor-bound conformations (closed, PDB:4AKE)^27^. It has been experimentally shown that at 35 °C the ATP_lid_ region unfolds selectively. This state is denoted as “binding incompetent” state and appears to be an intermediate state on the global folding pathway. Recently, Olsson et al.^30^ experimentally confirmed that the interconversion between open and closed state involves partial folding/unfolding of the ATP_lid_ subdomain.

In addition, the ATP_lid_ and AMP_lid_ conformational changes occur in absence of substrates^31, 32^.

In the AKe structure prediction the main challenge comes from the lack of small molecules that lock the protein in either the open or closed conformation. We selected the AK sequence range for which we have crystallographic information for both the apo (open) and the holo (close) conformations and generated 1000 models with our pipeline (see Method section for details). All the models were rescored with AWSEM force-field^33^; we have filtered out models with energies higher than -788 kcal/mol (see S1.A). The 189 remaining models were clustered based on their backbone RMSD for a total of two clusters (Figure 1.A). For each of the clusters a centroid has been selected as the one with the lowest energy (cluster 1: -1037.9 kcal/mol; cluster 2: -924.0 kcal/mol). By visually comparing the cluster’s population with the atomistic MD structure ensemble we initially observed that the conformational space explored by our models resembles the conformational shift between the apo and holo forms observed during the atomistic MD simulation (S1.B and S1.C). It’s interesting to notice that our models also explore an amplitude range not observed in the MD simulation. The highest flexibility of AMP_lid_ and ATP_lid_ regions is observed in both clusters as highlighted by the Root Mean Square Fluctuation (RMSF) analysis reported in Figure 1.B. The displacement intensity in the AMP_lid_ for cluster-2 (green) is roughly double (≈ 0.5 nm) compared to the one observed for cluster-1 (orange; ≈ 0.2 nm) and the 4AKE-MD (gray; ≈ 0.3 nm). Regarding the ATP_lid_ fluctuation, a similar displacement intensity is observed in both clusters (≈ 0.7 nm) which is roughly double the one observed in 4AKE-MD (≈ 0.4 nm). Principal Component Analysis (PCA) performed on cluster-1 and cluster-2 showed that the direction of the first principal component (PC1) involves the typical rearrangement of ATP_lid_ and AMP _lid_ regions experimentally reported^34^ and confirmed in the PCA performed on 4AKE-MD simulation frames (Figure 1.C). The PC1 density distribution (Figure 1.D) of cluster-1 (orange) exhibited a distribution mode compatible with the (i) open and close AK_e_ conformations and (ii) intermediate state conformations observed in the 4AKE-MD simulation (gray). Cluster-2 PC1 density distribution range (green) included the AK_e_ open conformation. By correlating the aperture angle with the distance between ATP_lid_ and AMP _lid_ regions we observed that cluster-1 (orange) and cluster-2 (green) explored both the close and open conformations respectively (Figure 1.E). The intermediate modes observed during the 4AKE-MD simulation (gray) were recapitulated by our models as well. RMSD clustering analysis on the currently available experimental structures deposited in the PDB showed that 35% of the adenylate kinase structures have an open-like conformation while the remaining 65% displayed a close-like conformation. The model clustering distribution is in agreement with the current experimentally available conformations showing 15% open-like models and 85% close-like models. AlphaFold2 models from ColabFold pipeline^25^ (red sphere) reviewed close and intermediate AKe conformations, while the AlphaFold2 prediction deposited in the AlphaFold Protein Structure Database (red star)^21^ provides only the close AKe conformation.

**Figure 1.**
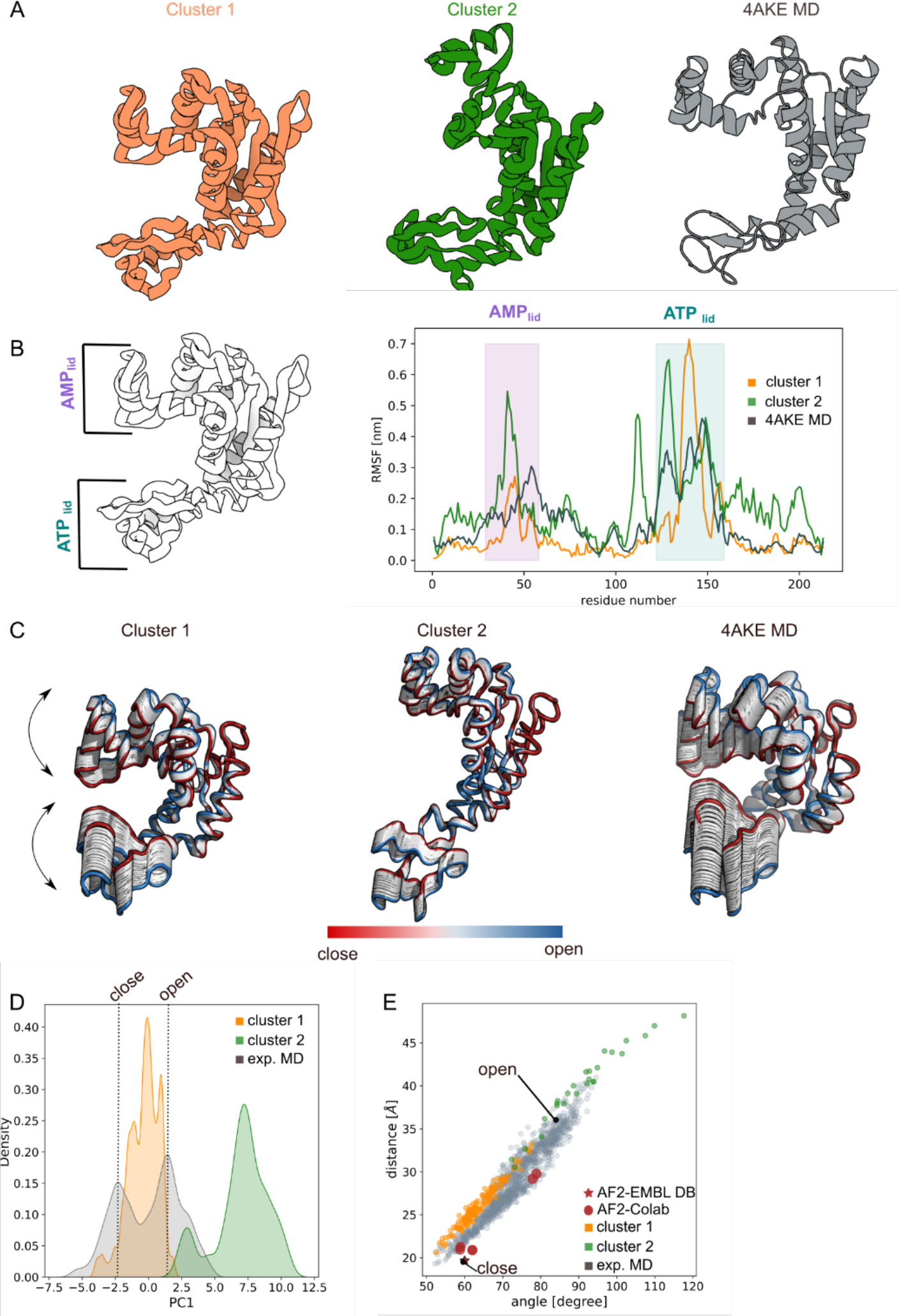
Adenylate kinase. **A.** Cartoon representation of the cluster centroids (orange and green) and 4AKE-MD snapshot (grey); **B.** Root Mean Square Fluctuation (RMSF) analysis showed a cluster flexibility in the ATP_lid_ and AMP_lid_ regions comparable to the one observed in the 4AKE-MD simulation; **C.** Principal Component Analysis for both clusters and MD simulations highlighted the protein internal movement showing for cluster-2 wider conformations than 4AKE-MD simulation; **D.** First principal component density distribution plot for 4AKE-MD (grey), cluster-1 (orange) and cluster-2 (green) unveiled that the cluster internal movement explored is compatible with the 4AKE-MD simulation; **E.** distance/angle correlation analysis of cluster-1 (orange) and cluster-2 (green) overlap with the conformational space explored by the 4AKE-MD closed and open conformations (gray). AlphaFold2 prediction from EMBLDB (red star) and Google Colab (red spheres) are also reported. The closed and open X-ray adenylate kinase conformation are specified with a black dot.

### **α**I-domains of LFA-1

Lymphocyte function-associated antigen 1 (LFA-1, integrin αLβ2) is a metal binding seven domain protein composed of a head piece (β-propeller, αI and βI) and two leg parts (β-propeller, Thigh, Calf-1/2, I-like domain, Hybrid Domain, I-EGF’s, β-tail). It belongs to the Integrin family of cell surface receptors whose functions are associated with leukocyte diapedesis, migration within tissue and the cell-adhesion recognition process. Precisely, LFA-1 binding to intercellular adhesion molecule 1 (ICAM-1) mediates the adhesion of leukocytes to blood vessels or antigen presenting cells (APC)^35–38^. Many integrins do not normally exhibit high affinity for ligands and must be activated in order to observe the binding^39–41^. This has led to the hypothesis that the change into the high affinity state is resulted in a conformational change. In response to several mechanical^42, 43^ and biochemical signals^36, 37, 44^, integrins undergo global and local conformational changes and ligand binding affinity variations. Under physiological conditions, LFA-1 ectodomains may assume a global bent conformation with a low binding affinity (PDB:5ES4^45^). Changing the metal ion from Ca^2+^/Mg^2+^ to Mn^2+^ results in an extended integrin conformation accompanied by a higher ligand binding affinity^36, 37, 46^ (PDB:5E6U^45^). In addition, local conformational changes occur at the level of the αI domain influencing the ligand binding affinity of LFA-1^38, 46^. The αI domain is the ligand binding domain of LFA-1 with a gradually decreasing affinity for metal ions in the order Mn^2+^ > Mg^2+^ > Ca^2+^ ^46, 47^. Indeed, structural studies on LFA-1 revealed a metal ion binding site called Metal Ion^48^ Dependent Adhesion Site (MIDAS) located at the top of the αI domain [9,32] ^49–54^. The MIDAS directly coordinates the sidechains of polar/acidic residues (S139, T206 and D239) discriminating between open and close conformations which correspond to high and low binding affinities respectively ^55–58^. In both open and close conformations the MIDAS ion shared S139 as primary coordinator. In the closed structure,T206 and D239 form direct bonds to MIDAS metal ion^51, 52^ while in the open conformations the metal ion is directly coordinated by T206 and the MIDAS ion undergoes inward movement by about 0.2 nm^59^. Surface plasmon resonance experiments suggest an inverse relationship between αI ligand affinities and metal binding: the closest conformation corresponding to the highest-ligand affinity^47^. Steering Molecular Dynamics simulation in explicit solvent confirms that the position of the metal ion is coupled with the α7-helix location (S283-T300): when the α7-helix is in the middle down position, the MIDAS ion has a strong tendency to move inward to its open position binding ligands with high affinity^59^. Indeed structural analysis on the αI domains^49–54^ shows that in the open conformation the metal ion coordination translates in a downward displacement of the α7-helix highlighting the presence of a transmission signal from the MIDAS of αI to the other integrin domains^60–62^. Lastly, the key residues in the “rachet ”-like αI region (L289, F292 and L295) have also been observed to be correlated with the movement of the α7-helix.

In this study we focused on the αI domain of the LFA-1 protein. The main challenge in the structure prediction of the αI/LFA-1 protein comes from the absence of the metal ion located at the MIDAS ion site which position is tied with the α7-helix location. All the MD simulations on the X-ray structures have been performed without the metal ions.

Two X-ray structures have been chosen as representative for the closed and open αI conformations: 1LFA^51^ and 3HI6^34^ respectively. We selected the αI sequence for which we have crystallographic information for both the open and close conformations and generated 1000 models with our pipeline (see Method section for details). All the models were rescored with AWSEM force-field^33^ and the ones with energy higher than -813 kcal/mol (see S1) filtered out. The 212 remaining models were clustered based on their backbone RMSD for a total of one cluster (Figure 2.A) while the centroid has been selected as the one with the lowest energy (cluster-1: -1038.5 kcal/mol). Visual inspection of cluster-1 revealed that our models sampled exclusively the closed conformation represented by 1LFA. The RMSD plot of the atomistic MD simulation on 1LFA and 3HI6 showed the stability of the initial X-ray structure (S2.B) confirming the inability to observe the transition between open and closed conformation without specific computational approach as SMD^63^. RMSF analysis (Figure 2.B) confirmed the low flexibility of the α7-helix (S283-T300; blue) for both cluster-1 (orange) and the 1LFA-MD simulation (dark gray). The difference in displacement between cluster-1/α7-helix (orange) and 3HI6-MD/α7-helix (light gray) is roughly 0.2 nm as observed between the crystal structures 1LFA and 3HI6. The “rachet ”-like αI movements in cluster-1 resembles the residue movement observed in 1LFA with the only exception of the cluster-1/F292 residue that showed higher displacement. The MIDAS residues flexibility (green) of cluster-1 appeared to be higher than 1LFA for residues S139 and T206 reflecting the lower sensitivity of Ca^2+^, Mg^2+^ and Mn^2+^ to the position amino acid conservation estimation^64^ and therefore in the resulting model prediction.

**Figure 2.**
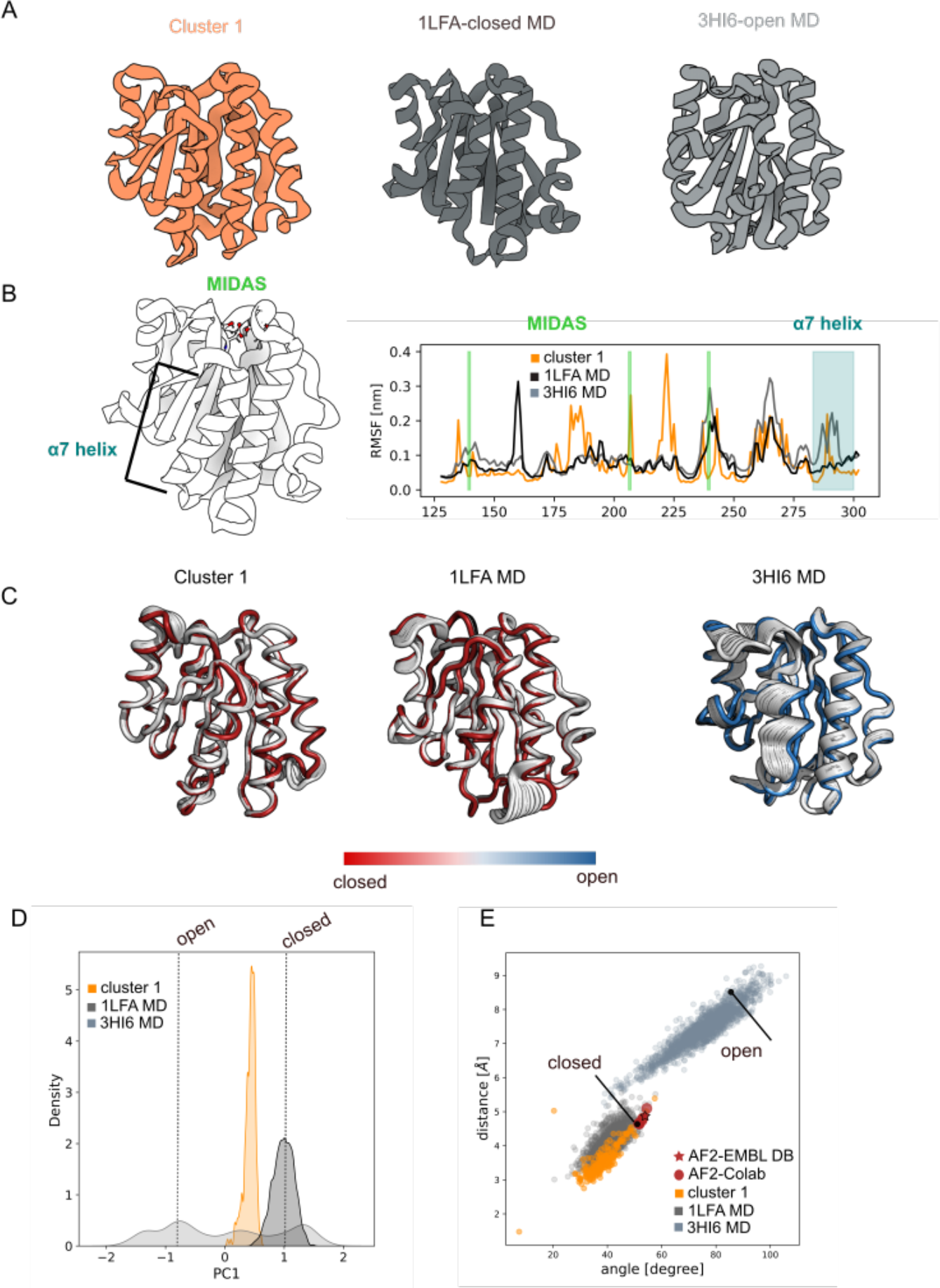
αI-domains of LFA-1. **A.** Cartoon representation of the cluster centroid (orange), 1LFA-MD snapshot (dark-grey) and 3HI6-MD snapshot (light-gray); **B.** Root Mean Square Fluctuation (RMSF) analysis showed the cluster rigidity α7-helix regions (orange) comparable to the closed 1LFA-MD conformation (dark-grey); **C.** Principal Component Analysis for the cluster and MD simulations highlighted the protein internal movement showing the cluster low flexibility compatible with the 1LFA-MD simulation; **D.** First component density distribution plot for 1LFA-MD (dark-grey), 3HI6-MD (light-gray) and cluster-1 (orange) unveiled that the cluster internal movement explored is compatible with the 1LFA-MD simulation; **E.** distance/angle correlation analysis of cluster-1 (orange) overlaps with the conformational space explored by the 1LFA-MD closed conformations (dark-gray), while none of our predictions explored the open 3HI6 conformation. AlphaFold2 prediction from EMBLDB (red star) and Google Colab (red spheres) are also reported showing conformations near the closed X-ray structure. The closed and open X-ray αI-domains of LFA-1 conformations are specified with a black dot.

Principal Component Analysis (PCA) performed on cluster-1 highlighted the stability of both α7-helix and MIDAS residues confirming the adoption of a close LFA-1 conformation rather than the flexible open conformations (Figure 2.C). The PC1 density distribution (Figure 2.D) of cluster-1 (orange) exhibited a distribution mode compatible with the close LFA-1 conformations observed in the 1LFA-MD simulation (dark gray). The correlation analysis between the angle and the distance of the α7-helix with respect to the body of the LFA1 protein confirmed that both our models (orange) and the AlphaFold2 models from ColabFold pipeline^25^(red dot) resembled a close LFA1 conformation (Figure 2.E). The intermediate modes observed during the 1LFA-MD simulation (dark gray) were recapitulated by our models as well, while the AlphaFold2 prediction deposited in the AlphaFold Protein Structure Database (red star)^21^ provides only the closed LFA1 conformation. RMSD clustering analysis on the current available experimental structures deposited in the PDB showed that 6% of the αI/LFA-1 structures have an open-like conformations and 79% displayed a close-like conformations. Interesting to notice 15% of the total experimental αI/LFA-1 structures have an intermediate-like conformations characterized by a displacement of the α7-helix position. The model clustering distribution captured only the most abundant experimental conformations correspondent to the closed-like α7-helix positions.

### Myoglobin protein

Myoglobin is a heme-containing globular protein that is abundant in myocyte cells of heart and skeletal muscle^65^. Its function involves (i) the reversible binding of oxygen to facilitate its diffusion^66^ and (ii) the scavenging of nitric oxide to prevent the inhibition of the mitochondrial enzyme cytochrome C-oxidase^67, 68^. Myoglobin has a molecular mass of roughly 17 kDa and consists of a single polypeptide of 153 amino acids with a secondary structure of eight α-helices. Between the fifth and the sixth α-helix there is a hydrophobic region that accommodates the heme moiety.

Several crystallographic structures have been solved for both the apo- and holo- Myoglobin^69–71^ showing that the main variation is due to the location of the α6-helix (from H82 to K98) where the heme pocket is located. The heme is bound to H95 providing extra stabilization and steric bulk that prevents collapse^72^. Previous protein structure prediction of the Myoglobin conformation showed that the absence of the heme made it impossible to provide stabilization and leading to greater structural heterogeneity of the α6-helix^73^.

We considered two forms of Myoglobin, the apo structure without its heme cofactor (PDB: 2EB8)^74^, and the holo form containing the cofactor (PDB: 2JHO)^75^. The overlap between the two X-ray structures showed a deviation in the α6-helix region of 4.3 Å: in the apo-form the α-helix is unfolded occupying the heme-pocket and coordinating the copper complex (Cu^II^(sal-X)), while in the holo-form the α6-helix is folded establishing the side of the heme pocket. The main challenge in the structure prediction of Myoglobin comes from the absence of both the heme cofactor and the metal coordinator. Our atomistic MD simulations on the X-ray structures have been performed without the heme and the copper complex Cu^II^(sal-X) showing that the α6-helix of the apo conformation assumed the same location of the α6-helix of the holo conformation during the first 10 ns of simulation (S3.A). Indeed, backbone RMSD analysis on the MD simulations revealed an increment of a factor of 2 Å for 2EB8 confirming the structural deviation from the initial X-ray position ( S3.B).

The same sequence coverage for both the apo- and holo-conformation was selected as input for our pipeline to generate up to 1000 models (see Method section for details). All the models were rescored with AWSEM force-field^33^ and models with energy higher than -664 kcal/mol were filtered out. The 212 remaining models were clustered based on their backbone RMSD for a total of one cluster (Figure 3.A) while the centroid was selected as the one with the lowest energy (cluster-1: -804.7 kcal/mol). Visual inspection of cluster-1 revealed a folded α6-helix which explores a similar location as the one observed on the MD simulation of the holo-structure (2JHO; Figure 3.B ). RMSF analysis (Figure 3.C ) showed a lower α6-helix residue flexibility for cluster-1 (orange) as compared to the holo- (dark gray) and apo-conformations (light gray) with main differences in the region between L40 and D60. PC1 eigenvector representation in Figure 3.C displayed less movement in the α6-helix region confirming the overall helical rigidity of the predicted models in contrast with the higher flexibility observed for the apo 2EB8-MD. The PC1 density distribution (Figure 3.D) of cluster-1 (orange) exhibited a distribution mode compatible with the holo-conformation partially sampling intermediate mode states between the apo- and holo-conformation. By correlating the aperture angle with the distance between α6- and α5-helix as heme pocket descriptors (Figure 3.E; see Supplementary Information) we observed an overlap between the conformational space explored by holo- (dark gray) and apo-conformation (light gray) showing the transition of apo-2EB8 structure to a 2JHO-like structure in the absence of metal coordinator (Cu^II^(sal-X)). This conformational conversion revealed that in the absence of any metal complex only one heme pocket arrangement is observed, which translates into one models. Cluster-1 (orange) explored the conformational space observed in the MD simulations (light and dark gray). RMSD clustering analysis on the current available experimental structures deposited in the PDB showed that 99% of the myoglobin structures have a holo-like conformation and 1% displayed an apo-like conformation. The α6-loop arrangement is observed only in two PDB structures (2EB8 and 2EB9). For this example, we were able to compare our results only with the ColabFold pipeline models ^25^(red dot) since the no prediction is available for myoglobinin the AlphaFold Protein Structure Database. ColabFold pipeline^25^ predictions explored only a limited region of the conformational space covered by the MD simulations in between the X-ray structures. It is worth mentioning that from this analysis we observed that the two clusters obtained are not representative of the two experimental conformations but reflect the flexibility observed in the region between L40 and D60.

**Figure 3.**
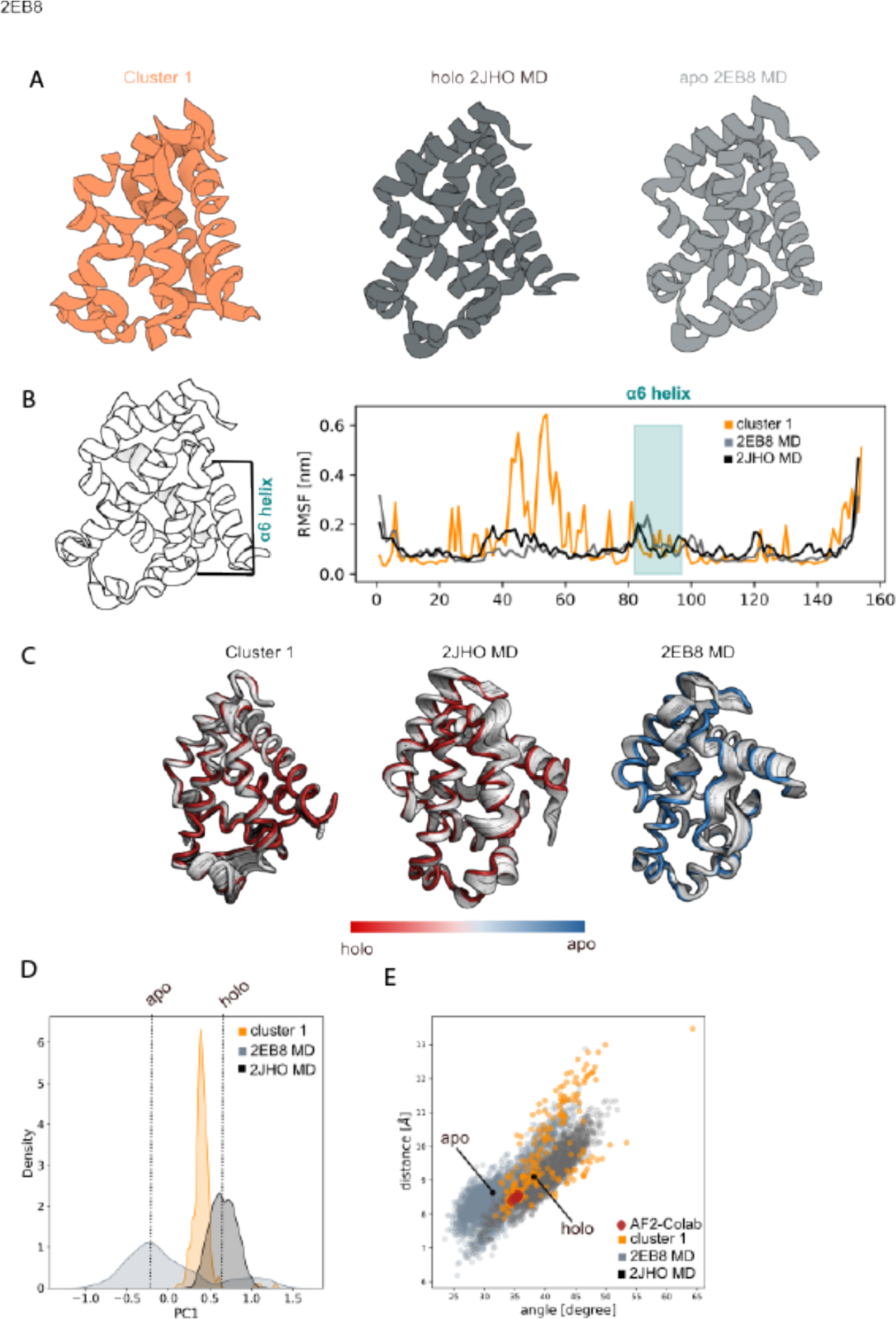
Myoglobin protein. **A.** Cartoon representation of the cluster centroid (orange), 2JH0-MD snapshot (dark-grey) and 2EB8-MD snapshot (light-gray); **B.** Root Mean Square Fluctuation (RMSF) analysis showed the cluster rigidity α6-helix regions (orange) comparable to the holo 2JH0-MD conformation (dark-grey); **C.** Principal Component Analysis for the cluster and MD simulations highlighted the protein internal movement showing the cluster low flexibility compatible with the 2JH0-MD simulation; **D.** First principal component density distribution plot for 2JH0-MD (dark-grey), 2EB8-MD (light- gray) and cluster-1 (orange) unveiled that the cluster internal movement explored is compatible with the 2JH0-MD simulation; **E.** distance/angle correlation analysis of cluster-1 (orange) overlaps with the conformational space explored by both the holo 2JH0-MD and apo 2EB8-MD (dark-gray). AlphaFold2 prediction from EMBL-DB was not public available. Google Colab predictions are reported as red spheres and explored intermediate states. The closed and open X-ray αI-domains of LFA-1 conformations are specified with a black dot.

### T4 lysozyme

T4 lysozyme (T4L) is a globular protein of roughly 18 kDa that helps release mature phages by breaking down the Escherichia coli cell wall during infection^76^. The three-dimensional structure is organized in two domains joined by a long helix: the N-terminal (residue 12-60) and the C-terminal (residue 80-164). The active site cleft is located at the interface between the two domains. An occluded active site were observed in early X-ray structures suggesting the hypothesis that relative movements between the N-and C-terminal domains enable substrate access^77^. Later crystal structures of point mutants (e.g. M6I) showed conformations where the active site is more open, suggesting that the two domains undergo a ‘hinge-bending’ motion^78, 79^. Direct evidence of the hinge-bending transformation was obtained by Electron Magnetic Resonance (EPR) analysis showing an equilibrium between open and close conformation in solution^80^. EPR distance analysis confirmed that the open structures are not purely distortions caused by the mutations and crystal packing but represent the consequence of intrinsic hinge-bending motions. This motion was furthermore supported by NMR^76^ data and MD^81, 82^ simulations, corroborating the presence of ‘hinge-bending’ motion in T4 lysozyme. Moreover, Fluorescence Correlation Microscopy (FSC) established the presence of multiple intermediate conformations, in contrast to the two state model^83^.

We considered two X-ray structures representing the open (PDB:150L) and close (PDB:256L) T4-lysozyme conformations. The main challenge in the structure prediction of the T4-lysozyme comes from considering the mutated protein sequence M6I as starting point of our prediction to test the ability of our pipeline in predicting conformational changes in the presence of point mutation.

The same sequence coverage for both the open- and close-conformation was selected as input for our pipeline to generate up to 1000 models (see Method section for details). All models were rescored with AWSEM force-field^33^ and the ones with energy higher than -537 kcal/mol (S4.A) filtered out. The 182 remaining models were clustered based on their backbone RMSD for a total of two clusters (Figure 4.A) and the centroid selected as the one with the lowest energy (cluster-1:-699 kcal/mol; cluster-2:-627 kcal/mol). Initial visual inspection of the two clusters showed a representative open and close conformation for cluster-1 and cluster-2 respectively. RMSF analysis (Figure 4.B) displayed high flexibility in correspondence of the N- and C-terminal domains. In particular, we noticed higher flexibility for both domains in 150L-MD simulation (dark gray) while lower flexibility for C-terminal domain in 256L-MD simulation (light gray). A similar behavior is observed for the clusters: (i) cluster-1 (orange) showed high flexibility in both the protein domains as for 256L-MD and cluster-2 (green) exhibited higher rigidity in the C-terminal region as for 150L-MD. Analysis on the PC1 eigenvector projection (Figure 4.C) revealed higher N- and C-terminal distance for cluster-2 as compared to cluster-1 while 256L PC1 seems to keep a closer conformation during the MD simulation. Further analysis on the PC1 density distribution (Figure 4.D) showed a tendency for cluster-1 to explore modes nearer the closer conformation while for cluster-2 to explore modes closer to the open conformations. The correlation plot between the distance and the angle formed by N- and C-terminal (Figure 4.E) confirmed the open conformational sampling for cluster-2 (green) and closed and intermediate conformational sampling for cluster-1 (orange). During the MD simulation 256L kept a closed conformation while 150L showed the typical ‘hinge-bending’ motion discussed above (S4.B). RMSD clustering analysis on the current available experimental structures deposited in the PDB showed that 21% of the T4 lysozyme structures have an open-like conformation while the remaining 79% displayed a close-like conformations. The model clustering distribution is in agreement with the current experimentally available conformations showing 11% for the open-like models and 89% for the close-like models. For this example, we were able to compare our results only with the ColabFold pipeline models ^25^(red dot) since the AlphaFold Protein Structure Database prediction is not available. ColabFold pipeline^25^ predictions explore only the closed region of the conformational space covered by the 256L-MD simulations.

**Figure 4.**
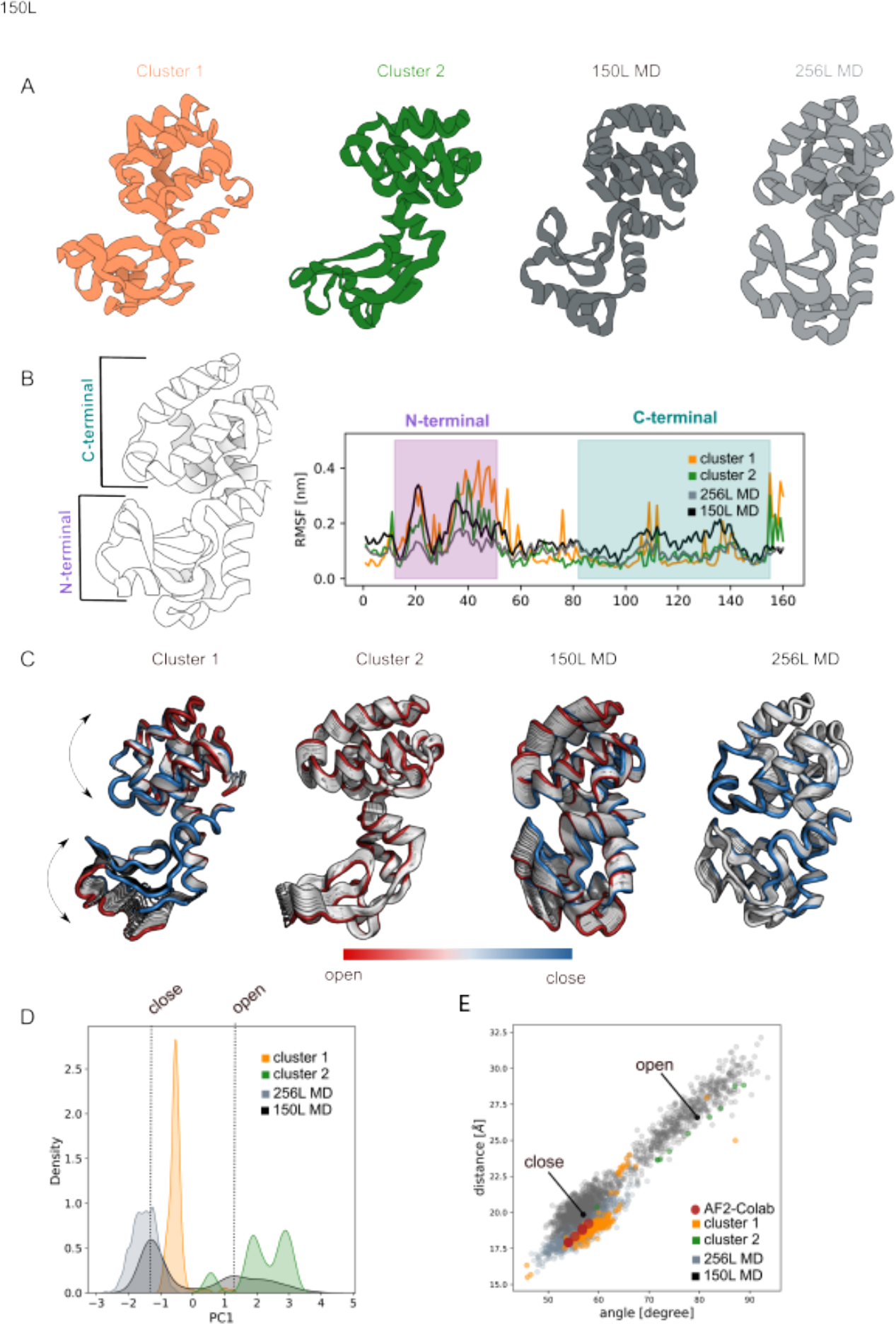
T4 lysozyme. **A.** Cartoon representation of the cluster centroids (orange and green), 150L-MD snapshot (dark-grey) and 256L-MD snapshot ; **B.** Root Mean Square Fluctuation (RMSF) analysis shows high flexibility in both the region for cluster-1 (orange) as observed for 150L-MD (dark-gray), while cluster- 2 exhibited low flexibility in the correspondence of C-terminal as observed for 256L-MD (light-gray); **C.** Principal Component Analysis for both clusters and MD simulations highlighted the protein internal movement showing a cluster-1 (orange) flexibility comparable to 150L-MD (dark-gray) and cluster-2 (green) flexibility comparable to 256L-MD (light gray); **D.** First principal component density distribution plot for 150L-MD (dark-grey), 256L-MD (light-grey), cluster-1 (orange) and cluster-2 (green) unveiled that the clustered-1 and clustered-2 models resemble the closed and open conformation respectively; **E.** distance/angle correlation analysis of cluster-1 (orange) and cluster-2 (green) overlap with the conformational space explored by the closed 150L-MD and open 256L-MD conformation respectively. AlphaFold2 prediction from EMBLDB was not public available. Google Colab model prediction (red spheres) explored closed conformation space. The closed and open X-ray T4 lysozyme conformations are specified with a black dot.

### Tetrahymena thermophila-BIL2

BIL2 is a part of the polyubiquitin locus of *Tetrahymena thermophila* (BUBL) where two bacterial-intein like (BIL) domains are flanked by two independent ubiquitin like domains (ubl4*ubl5). Generally, inteins are protein sequences known to perform protein splicing producing an intein domain and an inteinless host protein without insertion^84^. On the contrary, the BIL2 domain does not disrupt the structural continuity of the protein but rather interposes between two independently folded domains^85^. Their ability of acting as promiscuous catalytic elements that exercise protein splicing between several different domains has been extensively used in protein engineering and synthetic biology^86–88^. From a biological function perspective, the intein protein-splicing translates into a benefit for the host by constituting a switch from inactive to active native protein. In particular, BIL2 presents another interesting aspect: the presence of a highly conserved motif, RGG, that represents the biological hallmark for ubiquitin activation and conjugation^89^. It has been experimentally observed that *T. thermophila* BIL domains are able to provide a perfect activated ubl protein without energy consumption^90^.

Recently, two structures of the splicing element TthBIL2 (PDBs:6TMM, 6Y75) have been deposited in the PDB shedding light on the protein splicing reaction and the role of the metal ion as reaction promoter rather than inhibitor^91–93^. The BIL2 displayed a well recognizable horseshoe-like fold typical for HINT domains, but in contrast to the majority of them, no α-helix structural elements are present. At the moment, only these two structures are publicly available representing both the inactive and unprecedent zinc-induced active forms. Structural comparison between the apo (PDB:6TMM) and the zinc-bound form (PDB:6Y75) allowed the elucidation of the role of zinc for protein splicing . The zinc is bound away from the catalytic center to which it is connected only through a H-bond network. It is coordinated by H48 and H125 disrupting the secondary structural arrangement between the two exteins. Based on the latest model, the zinc-binding regulates the splicing reaction by inhibiting the premature C-cleavage through the H125 interaction and by inducing a conformational change upon activation of N-S acyl shift which induces the thio-ester bond formation with the N-extein.

The same sequence coverage for both the apo- and holo-conformation was selected as input for our pipeline to generate up to 1000 models (see Method section for details). All the models were rescored with AWSEM force-field^33^ and the models with energy higher than -564 kcal/mol (S5.A) filtered out. The 179 remaining models were clustered based on their backbone RMSD for a total of one cluster (Figure 5.A) and the centroid selected as the one with the lowest energy (cluster-1:-659 kcal/mol). Initial visual inspection of the cluster (Figure 5.A) showed the same global folding of both the experimental structure, in agreement with the MD simulations of the X-ray structures (S5.B). RMSF analysis showed a cluster flexibility (orange) comparable with the 6TMM-MD (light-grey) and 6Y75-MD (dark-grey) with the exception of the region between D20 and I40 where higher movement is observed for the cluster. It’s important to mention that the flexibility of the key histidine residues (H48 and H125) in cluster-1 is in agreement with the experimental MD simulations (Figure 5.B). Analysis on the PC1 eigenvector projection (Figure 5.C) confirmed a correlation between the flexibility of cluster-1 with the 6Y75-MD and 6TMM-MD showing that the apo and Zn-bound conformations are sampled in all the structure ensemble. We noticed once more that the protein region between I40-Q60 present in cluster-1 is flexible as compared to the X-ray MD simulations (Figure 5.D). Detailed analysis on the H48 and H125 residue orientations revealed that both the experimental structures sampled the apo and Zn-bounded histidine residue orientations in the absence of the Zn^2+^ ion. Cluster-1 (orange) sampled as well an orientation space compatible with both the apo and Zn-bounded, while the five models generated by ColabFold pipeline^25^ explored orientation closer to the apo-form (red dots) (Figure 5.E). AlphaFold2 prediction from EMBL database provides a conformation closer to the Zn-bound conformation.

**Figure 5.**
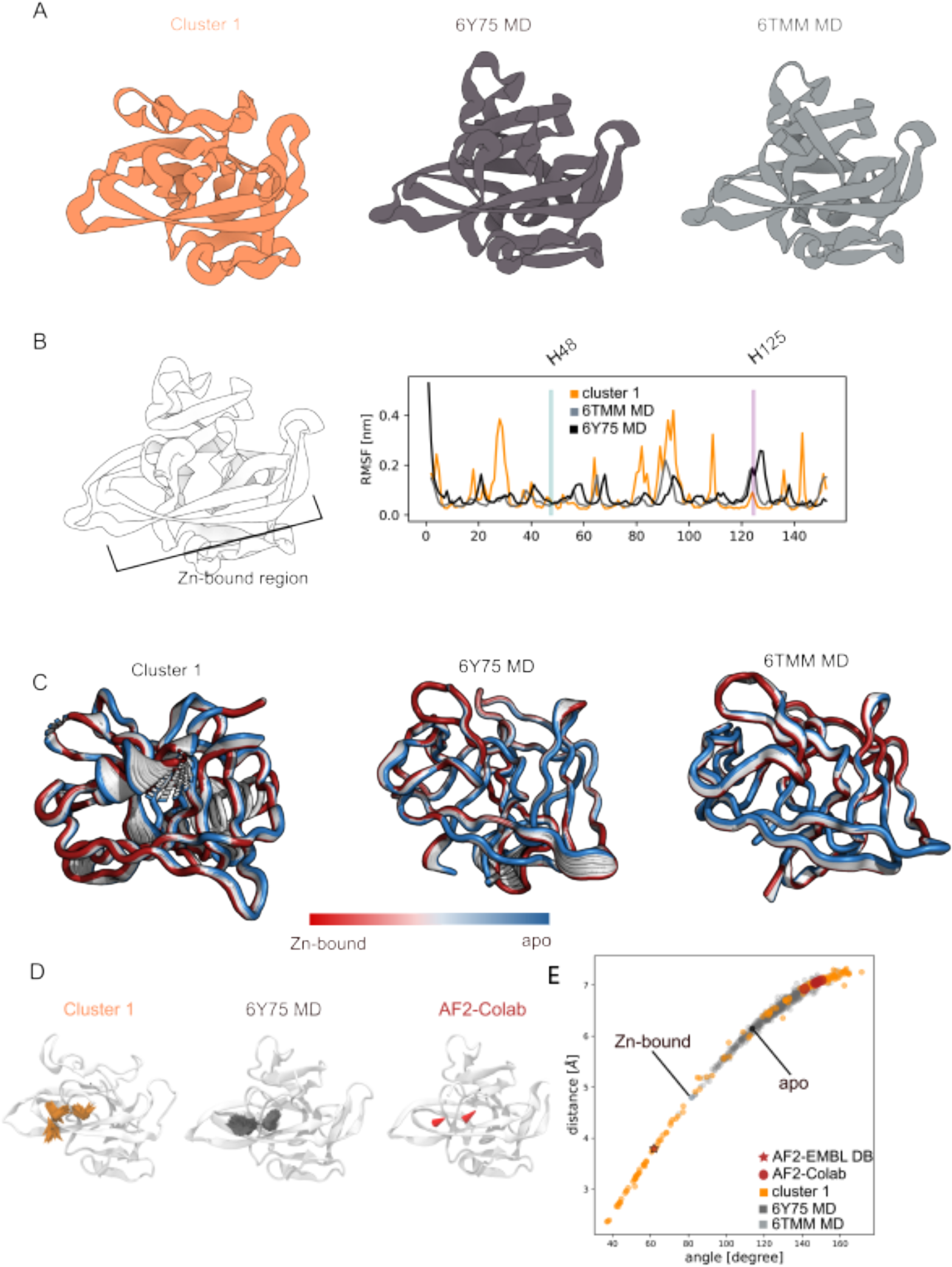
Tetrahymena thermophila-BIL2. **A.** Cartoon representation of the cluster centroid (orange), 6Y75-MD snapshot (dark-grey) and 6TMM-MD snapshot ; **B.** Root Mean Square Fluctuation (RMSF) analysis shows H48 rigidity for both the cluster (orange) and the experimental structures (6Y75 in dark grey; 6TMM in light-gray). The H125 flexibility is lower for cluster-1 compare to 6Y75 and 6TMM; **C.** Principal Component Analysis for cluster-1 and MD simulations highlighted the protein internal movement showing a cluster-1 (orange) flexibility comparable to both 6Y75-MD (dark-gray) and 6TMM-MD (light gray); **D.** Graphical representation of the H48 and H125 orientations for cluster-1 (orange); 6Y75-MD (dark-grey) and Google Colab predictions (red); **E.** distance/angle correlation analysis of cluster-1 (orange) overlap with the conformational space explored by both the Zn-bounded 6Y75-MD and apo 6TMM-MD conformation respectively. AlphaFold2 prediction from EMBLDB was not public available. AlphaFold2 prediction from EMBLDB (red star) and Google Colab (red spheres) are also reported. The Zn-bounded and apo Tetrahymena thermophila-BIL2 conformations are specified with a black dot.

## Method

For each of the 5 test cases (4 from deposited PDB structures publicly available, 1 AZ internal PDB) the following pipeline has been applied to generate the initial coarse models (Figure 6)

**Figure 6.**
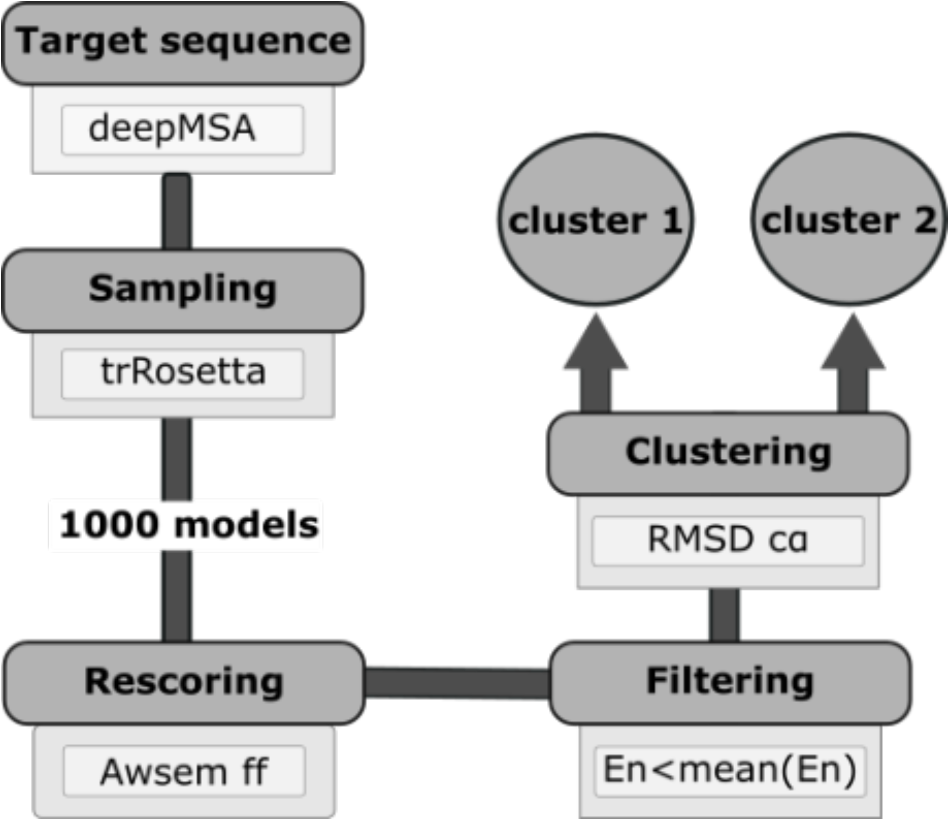
Flowchart of the AZ protein-folding pipeline. Four stages of the protein-folding prediction are performed consecutively: (1) MSA profile generation with DeepMSA, (2) protein model predictions with trRosetta, (3) energy rescoring with AWSEM Hamiltonian and energy filtering and (4) Hierarchical RMSD clustering with MDTraj.

### Multi Sequence Alignment generation

Multi Sequence Alignment (MSA) provides useful information about the evolutionary conserved positions and motifs that can hardly be derived from a single query sequence. In protein structure prediction, MSA is the principal source to retrieve local feature and residue-residue contacts which becomes critical for *ab initio* protein structure prediction ^94^.

Several sequence construction methods have been derived over the past years. PSI-BLAST is one of the most used approaches to query sequence specific profile generation^95^. HHblits^96^ from the HH-suite has recently become the general method for profile hidden Markov model (HMM) construction, and HMM search tools from HMMER suite^97^ are valid alternatives for the applications. However, only few pipelines efficiently provide sensitive MSA profiles from the query sequence by scanning multiple large sequence databases. DeepMSA^98^ is an open-source method for constructing sensitive and highly diverse MSA by merging sequence from three whole genome and metagenome databases. It combines HHblits and modified version of Jackhammer/HHsearch^99^ to perform homologous sequence search and further refine the alignment with a custom HHblits database reconstruction step. It has been proven that deepMSA consistently improves the accuracy of contact and secondary structure prediction which are particularly important for protein-structure prediction^98^.

We compared the trRosetta MSA profile generation with HHblits and deepMSA to ensure the accuracy of the MSA input. As a test case we considered the myoglobin protein (PDB: 2EB8) and generated the MSA profile with: (i) the default trRosetta MSA pipeline^24^ , (ii) the default deepMSA pipeline and (iii) HHblits by iteratively scanning three times (-n 3) the Uniref30 database^100^ for homologous sequence with a minimum master sequence coverage (-cov) of 75% and default e-value cutoff. The quality of the MSA profile was determined by comparing the probability distance prediction generated by trRosetta (see trRosetta section below) between two residues H98 and S93 which are observed to be at different distance in the open (d≈6 Å) and close (d≈8 Å) conformation. The distance prediction with trRosetta-MSA showed a shoulder at 7 Å and a pick at 9 Å (S6.A) while with deepMSA (S6.B) we have a separation between the pick at 6 Å and 8 Å. HHblits-MSA distance prediction showed a completely different shift in the trend with only one pick at roughly 8 Å. Due to the importance of the MSA quality in the final protein structure prediction we opted for deepMSA to generate the MSA profile input.

### trRosetta and the energy filtered step

1000 models have been generated with the XML version of transform-restrained Rosetta pipeline (trRosetta) used during CASP13 and available at https://github.com/gjoni/trRosetta24. The procedure adopted in our study does not involve the use of templates to test the ability of the deep residual network to predict structural information for proteins that lack structural homologues in the PDB.

The trRosetta deep neuronal convolutional network provides probability distributions for Cβ-Cβ distance and inter residue orientations from the MSA and converts them into spline restraints. Furthermore, the Rosetta model building minimization protocol guided by the restrains generates final predictions which are scored with the Rosetta score function ref2015^101^ as well as the Associative memory, Water mediated, Structure and Energy Model (AWSEM) force field^33^ . AWSEM is a coarse-grained protein force field used for *de novo* protein structure prediction which contains physical motivated terms and bioinformatically based local structure biasing terms that take into account many-body effects modulated by the local sequence. The comparison between the RMSD/Rosetta-Energy plot with the RMSD/Awsem plot (S8) showed a higher correlation for the latter consenting to filter out the models with energy higher than the mean Energy and proceed with the further analysis.

### RMSD clustering and centroid selection

The energy filtered models have been hierarchically clustered using pairwise RMSD implemented in MDAnalysis Tools^102^. The number of clusters that best represent the heterogeneity of the data set has been chosen with the Elbow method^103^. For each of the generated clusters a representative model is selected as the one with the lowest energy among all the other models in the cluster. Furthermore each of the lowest energy models per cluster is refined.

### Ensemble Analysis

For each of the test cases we performed a series of analysis to evaluate the conformational space explored by the models. The Principal Component Analysis (PCA) has been adopted to characterize the general folding features of proteins and the vector orientation analysis to investigate onto specific conformational transition of the target proteins. The visual representation and the density plot of the first principal component captures and describes in detail the overall dynamics of a system while relative residue fluctuations are examined by the Root Mean-Square Fluctuation (RMSF). We considered as ground truth the conformational space explored by the Molecular Dynamics simulation of the experimental X-ray structure.

### Molecular Dynamics protocol

The initial experimental structure is downloaded from the PDB and only the protein coordinates are considered for the simulation. The resulting cleaned-model was fully solvated with TIP3P water models^104^ in a water box of the dimension 80 x 80 x 80 Å and neutralized by the addition of NaCl at a concentration of 150 mM. Amber99SB force field^105^ was used for running the simulation. MD simulations were performed in GROMACS.2019 software^106^. The system was minimized for 1,000 steps and equilibrated in the isothermal-isobaric (NPT) ensemble for 2 ns at 1 atm and 300 K using a time step of 2fs. The system was simulated for on average 30 ns in the NPT ensemble.

## Discussion

### ML behavior in predicting protein conformational changes and current limitations

The current protein structure prediction methods have been tested on the ability to predict static X-ray structures as the best represented in the training set which is generally the most predominant conformation in the PDB. However, proteins in their biological environment exist in a dynamic ensemble of conformations distributed across a free energy landscape according to their Boltzmann-weighted probability of occurrence. Experimental methods as X-ray diffraction^107^, NMR^108^ and Föster resonance energy^109^ normally provide either the average position of a large number of structures or the distribution of small numbers of structural properties restricting the conformational protein space to a trivial number of conformational snapshots rarely obtained under biological conditions. Therefore, the necessity to investigate the computational ability in predicting the protein conformational space. The deep learning protein prediction algorithms provide models with a resolution comparable to the experimental techniques listed above starting a new era in the folding problem field. Recently, a correlation between co-evolutionary analysis and protein flexibility has been observed showing the possibility to predict protein conformations with the only knowledge from the 1^st^ protein sequence. Therefore, we decided to investigate the ability of the current deep learning algorithms to predict protein conformational space.

In this study we compared the performance of trRosetta with the current state-of-the-art method AlphaFold from ColabFold Google implementation^25^. The main difference between trRosetta^24^ and AlphaFold^21^ is the way the structural information is retrieved from the MSA profile: trRosetta explicitly calculates the distance probability from the MSA coevolutionary analysis, retrieves the potential restraints to reduce the degree of freedom during the minimization and proposes the final structure. Alternatively, AlphaFold enables end-to-end structure prediction for which no additional post-processing steps is required and the final result can be directly evaluated with the error passing back along the network. The conformational analysis performed on the five models generated by AlphaFold confirmed the high atomic accuracy in predicting models closer to the X-ray structure exploring a conformational space near one of the experimental conformations. On the other hand, the models generated with our pipeline covered a greater conformational area sampling also intermediate conformations observed in MD simulations. On average, the delta backbone RMSD covered by the AlphaFold models in all the test cases is smaller (ΔRMSD ≈ 2 Å) than the one covered by the trRosetta ensemble (ΔRMSD ≈ 7 Å) indicating the higher precision of AlphaFold in describing the experimental structure as well as the limited protein dynamic space explored. It is important to mention that with ColabFold implementation of AlphaFold we were only able to generate five models rather than the 1000 models generated with our pipeline. Further investigation on the influence of number of models onto the AlphaFold conformational exploration needs to be performed.

The depth of the initial MSA as well as the choice of the energy function represent key points in protein conformational prediction. We observed that the number of effective sequences considered and the choice of the approach used to generate the MSA profile are fundamentals in the ability of deep learning algorithms as trRosetta to retrieve flexible information. The accuracy of the energy functions is an essential component in computational protein structure prediction. Assuming that we have a good discrimination between folded and unfolded structures we should satisfy both low energy and low RMSD. The native state of the protein is expected to be found at the minimum of the energy function in order to discriminate between native and non-native structure. We adopted the AWSEM forcefield which has been trained to differentiate decoys and identify the near-native structure as the one with low energy. The correlation between AWSEM energy function and RMSD shows the ability of the forcefield to discriminate between conformations allowing to energy filter the models. The energy evaluation of the experimental structures showed the presence of decoys with energies comparable to the experimental structure for all the test cases except the αI-domains of LFA-1 where only the energy of the experimental apo structure is considered in the energy filtered ensemble.

A further limitation of the current deep learning techniques is the uncertainty in the prediction accuracy of the protein active site. AlphaFold and trRosetta have been trained with the apo protein structures and only consider X-ray structures limiting the data set to static information. We investigated the ability of these deep learning algorithms to predict the side chain orientations for the test case Tetrahymena thermophila-BIL2 which experimentally discriminates the holo- and apo-form on the basis of H48 and H125 orientations. Our pipeline was able to predict the presence of the only global fold experimentally observed as well as to sample the several histidine orientations. ColabFold predicted the apo conformation and AlphaFold the Zn-bound structure. However, we failed on clustering out the different orientations showing the limitations of the current pipeline in discriminating between side chain conformations. It is important to notice that if the conformational transition depends on the presence of coordinated metal ions, the prediction is challenging and only one conformation was generated by our pipeline as well as ColabFold platform.

## Conclusions

Proteins are not static entities but in motion in the cell. Switching between structurally distinct states is common in proteins carrying out functions such as catalysis or molecular recognition. X-ray crystallography provides a static snapshot of the possible conformations assumed by the protein in a crystallization setting that is not representative of the physiological environment. NMR structures offer dynamic information of the protein in solution but the protein size is a limit difficult to overcome. Cryo-EM delivers an overview of the protein orientations but current technical limitations (e.g low signal to noise ratio) hamper high-throughput experiments preventing high numbers of structures to be available at high resolution. Hence the necessity of providing new approaches that allow to accurately and robustly predict conformational space and motions that are critical for protein function.

Alternative protein states are still a largely open question in the field of protein folding. With the advent of machine learning techniques for protein structure predictions as AlphaFold or trRosetta we reach accuracies of the overall fold prediction comparable with the experimental structures provided in the PDB. However the quality of the prediction is still evaluated on the basis of static experimental structure. Here, we tested the ability of the current state-of-the-art machine learning methods in predicting protein conformational space. We have observed a potential correlation between residue-residue distance prediction and protein flexibility. MSA seems to encode information regarding the conformational flexibility of the protein which can in turn translate into bimodal behavior of the probability distance and angle distribution used in machine learning approaches as trRosetta. By combining trRosetta^24^ with deepMSA for MSA profile generation^98^ and AWSEM force field^33^ for rescoring the predicted models, we observed that the conformational space explored by the predicted models was similar to the one observed during the MD simulations of the target experimental protein structure. In particular, for the adenylate kinase and the T4 lysozyme predictions it was possible to predict the active and inactive structures as well as the intermediate conformations observed during the MD simulations. Although the trRosetta algorithm has been trained on static PDB structures, it was able to a certain extent to reproduce the protein flexibility.

Limitations of the technique have been observed in the case where metal ions (LFA1 test case) and cofactor (Myoglobin test case) were influencing the conformational equilibrium. The clear conformational distinction observed experimentally was also lost during the MD simulations without the protein counterpart. It is important to notice that the ability to predict protein flexibility is correlated to the number of available structures for the different protein conformations in the PDB, and this is the strongest limitation of the current machine learning techniques. Another important aspect is the quality of the initial MSA profile. We observed in our study that the choice of the MSA algorithm can hamper the model prediction by favoring one conformation (see Supplementary Material). Not only the MSA profile quality but also mutations in the initial target sequence could have an impact in the performance of the final prediction with the current deep learning methods. A recent study on the end-to-end RoseTTAFold protein prediction showed that small perturbation in the input sequence lead to radically different predicted protein in contrast with the observation that similar sequence lead to similar structure. By modifying 5 residues in the target sequence the RMSD was spanning from 0.9 A to 34.2 A. This highlighted the sensitivity of RoseTTAFold^110^ to the variation in MSA profile confirming the low correlation between MSA depth and model accuracy previous observed^110^. On the contrary, trRosetta showed higher correlation between MSA depth and model accuracy that could in principle be translated in higher robustness of the method and ability to predict relevant conformational change upon mutation.

With our study we only begin to explore the capability of deep learning methods in conformational protein structure prediction. We observed the potential to extrapolate flexibility from MSA that translates into restraints providing models with biologically relevant conformations. However, all these methods currently rely strongly on the current structures available in the PDB - limiting the prediction performance on novel fold and unseen conformations. Introducing dynamic information from FRET experiments, conformational heterogeneity data from cross-linking Mass Spectrometry as well as increased numbers of cryo-EM structures in the training data set would bring a new set of information that would enhance the description of protein flexibility. A complete and accurate description of the protein conformational space would impact drug discovery by providing initial starting points for active and inactive conformations changing the way to approach the drug discovery campaign and allowing to focus on biologically relevant questions rather than methodological issues.

## Acknowledgments

We thank Giuseppina La Sala for the fruitful discussion during the project and the article drafting. M.A. is a fellow of the AstraZeneca R&D postdoc program.

## Notes

The authors declare the following competing financial interest(s): All the authors are employees of AstraZeneca PLC. M.A, **W.C, L. D. M., H. K., G. P., L. T., C. T. and J. U.** are employees of AstraZeneca and may own stock or stock options.

