## Supplementary Information for "Protein structure dynamic prediction: a Machine Learning/Molecular Dynamic approach to investigate the protein conformational sampling"

**Support Information**

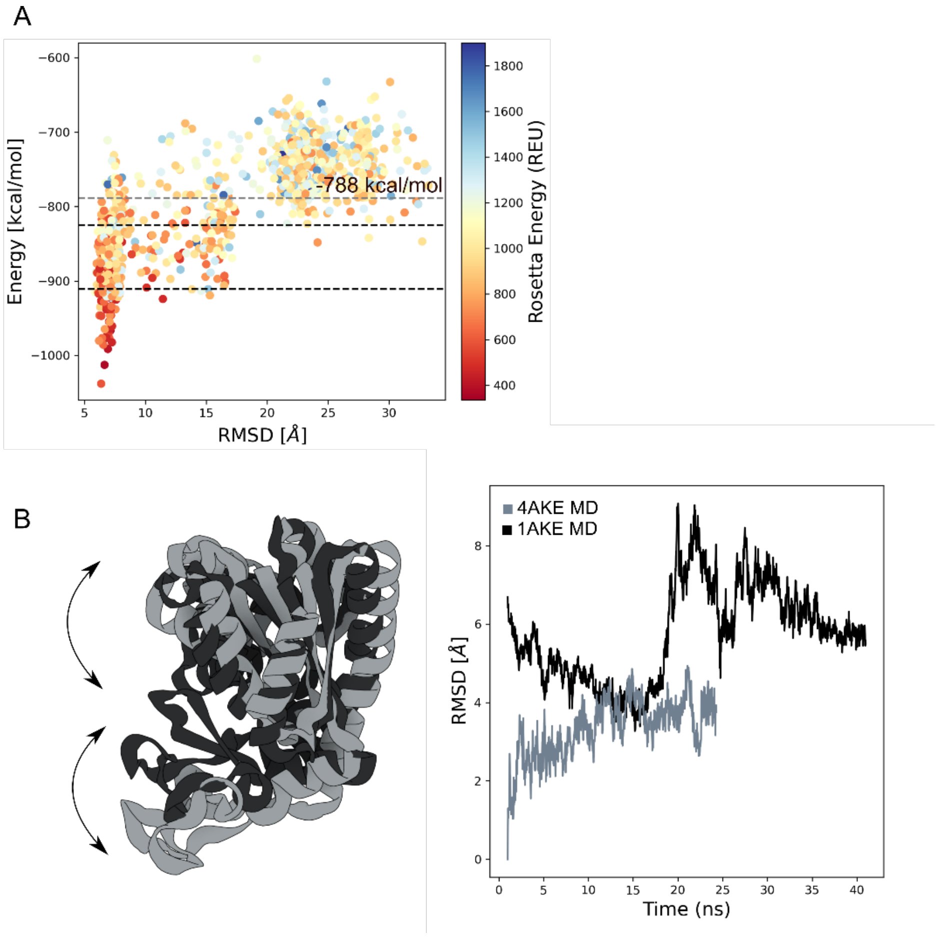


**Figure S1. Adenylate kinase A.** AWSEM force field vs RMSD profile plot. Each dot represents a protein prediction scored with AWSEM force-field and colored by the Rosetta energy score function ref15. The grey dashed line is the Energy mean filter. The black dashed line represent the X-ray structures energy value; **B.** left panel: overlap between the X-ray structure; right panel: Root-mean-square deviation (RMSD) plot of the 4AKE (light-grey) and 1AKE (dark-grey) protein during molecular dynamics simulation.

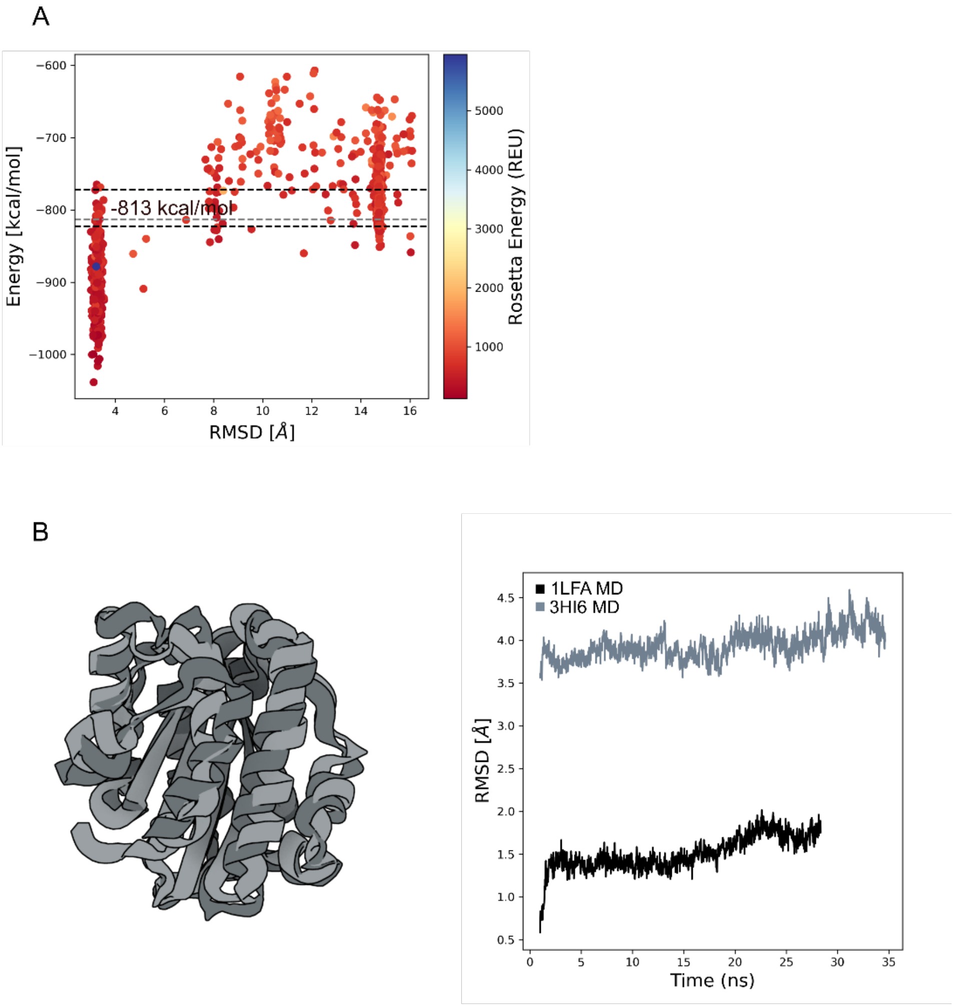


**Figure S2.** α**I-domains of LFA-1. A.** AWSEM force field vs RMSD profile plot. Each dot represents a protein prediction scored with AWSEM force-field and colored by the Rosetta energy score function ref15. The grey dashed line is the Energy mean filter. The black dashed line represent the X-ray structures energy value; **B.** left panel: overlap between the X-ray structure; right panel: Root-mean-square deviation (RMSD) plot of the 3HI6 (light-grey) and 1LFA (dark-grey) protein during molecular dynamics simulation.

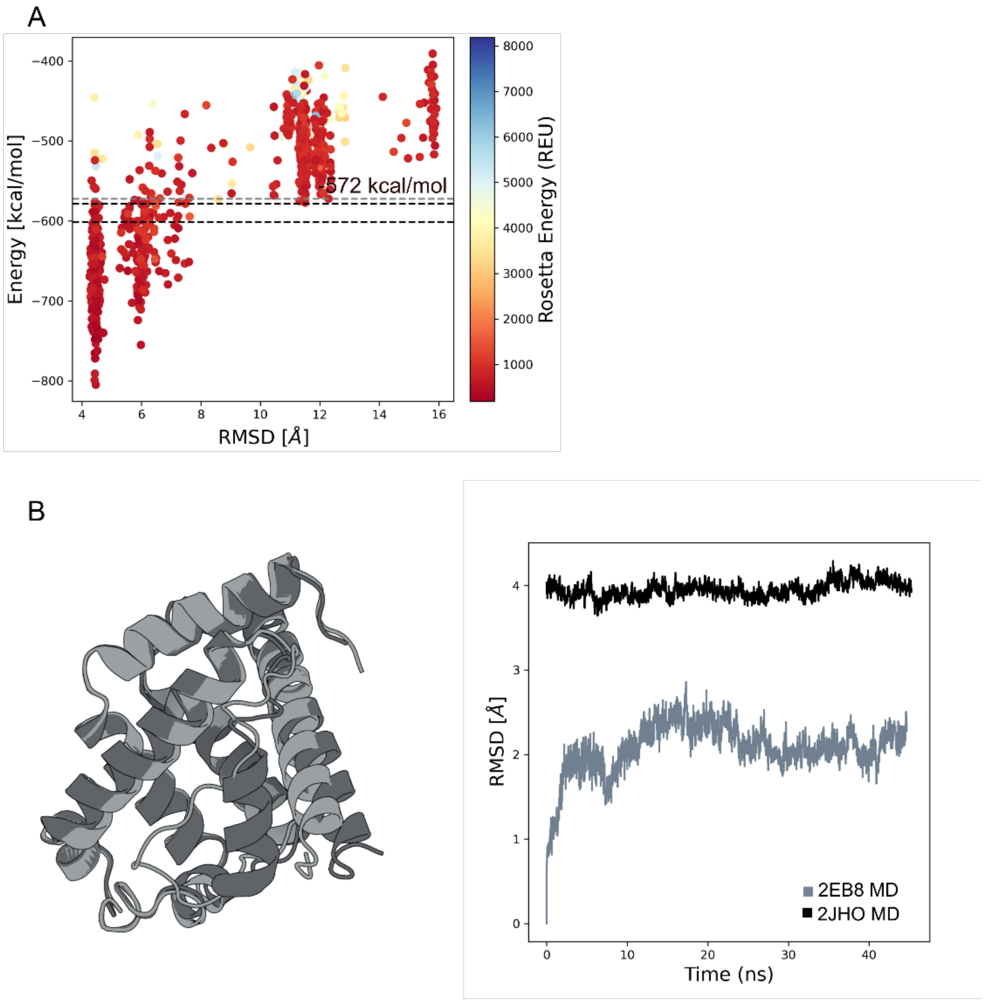


**Figure S3. Myoglobin protein. A.** AWSEM force field vs RMSD profile plot. Each dot represents a protein prediction scored with AWSEM force-field and colored by the Rosetta energy score function ref15. The grey dashed line is the Energy mean filter. The black dashed line represent the X-ray structures energy value; **B.** left panel: overlap between the X-ray structure; right panel: Root-mean-square deviation (RMSD) plot of the 2EB8 (light-grey) and 2JHO (dark-grey) protein during molecular dynamics simulation.

A

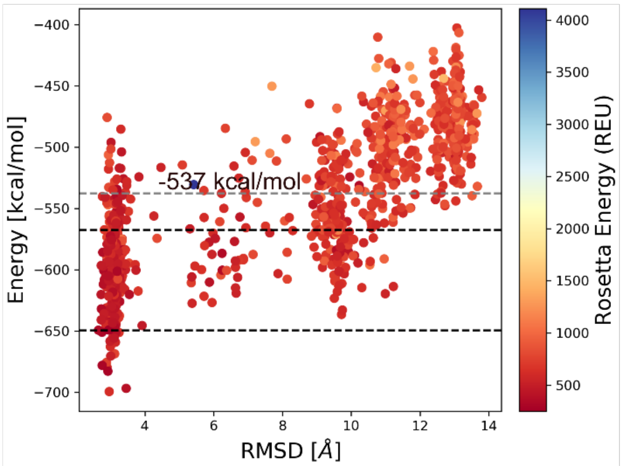


B

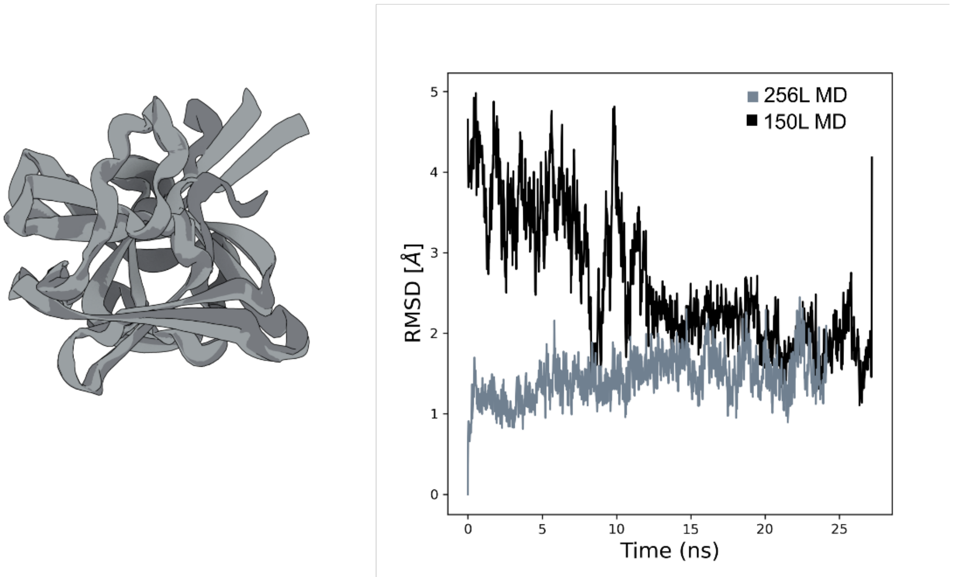


**Figure S4. T4 lysozyme. A.** AWSEM force field vs RMSD profile plot. Each dot represents a protein prediction scored with AWSEM force-field and colored by the Rosetta energy score function ref15. The grey dashed line is the Energy mean filter. The black dashed line represent the X-ray structures energy value; **B.** left panel: overlap between the X-ray structure; right panel: Root-mean-square deviation (RMSD) plot of the 256L (light-grey) and 150L (dark-grey) protein during molecular dynamics simulation.

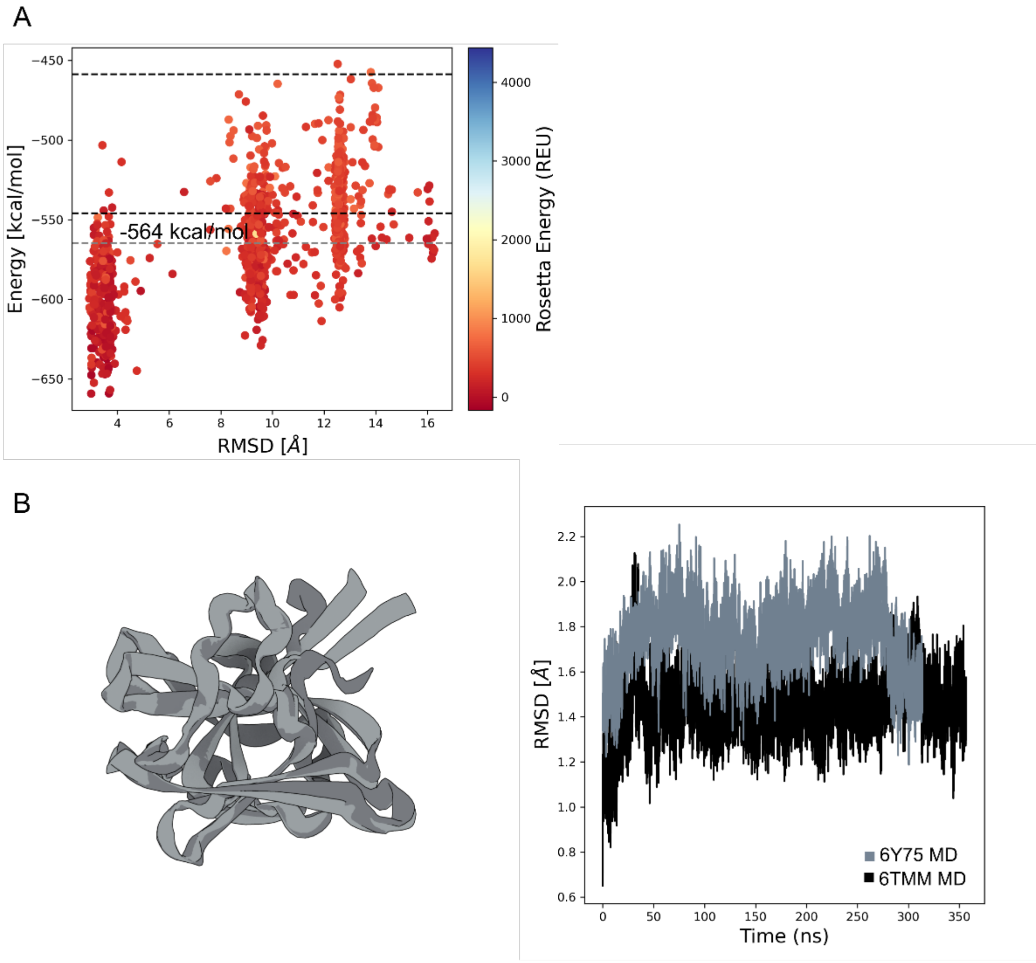


**Figure S5. Tetrahymena thermophila-BIL2. A.** AWSEM force field vs RMSD profile plot. Each dot represents a protein prediction scored with AWSEM force-field and colored by the Rosetta energy score function ref15. The grey dashed line is the Energy mean filter. The black dashed line represent the X-ray structures energy value; **B.** left panel: overlap between the X-ray structure; right panel: Root-mean-square deviation (RMSD) plot of the 6Y75 (light-grey) and 6TMM (dark-grey) protein during molecular dynamics simulation.

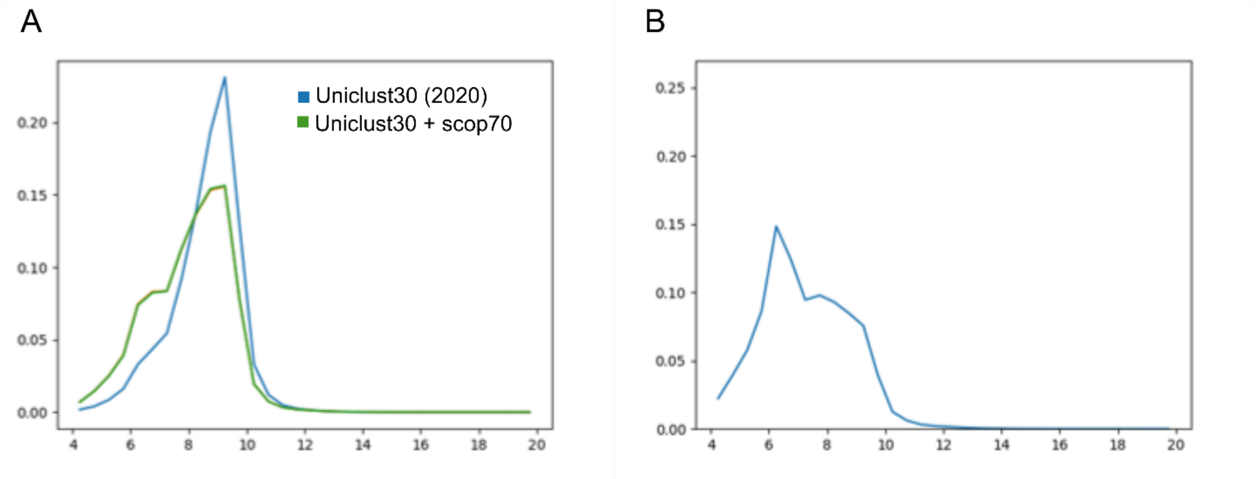


**Figure S6. Probability distance distribution: MSA effect.** Probability distance distribution between S93 and H98 calculated for the protein structure 2EB8. **A.** Probability distribution plot obtained with trRosetta-MSA protocol using Uniclust30 database (blue) and Uniclust30/scop70 (green). **B**. Probability distance distribution plot obtained with trRosetta-deepMSA protocol using Uniclust30 database (blue).
